# Improved Genomic Prediction Performance with Ensembles of Diverse Models

**DOI:** 10.1101/2024.09.06.611589

**Authors:** Shunichiro Tomura, Melanie J. Wilkinson, Mark Cooper, Owen Powell

## Abstract

The improvement of selection accuracy of genomic prediction is a key factor in accelerating genetic gain for crop breeding. Traditionally, efforts have focused on developing superior individual genomic prediction models. However, this approach has limitations due to the absence of a consistently “best” individual genomic prediction model, as suggested by the No Free Lunch Theorem. The No Free Lunch Theorem states that the performance of an individual prediction model is expected to be equivalent to the others when averaged across all prediction scenarios. To address this, we explored an alternative method: combining multiple genomic prediction models into an ensemble. The investigation of ensembles of prediction models is motivated by the Diversity Prediction Theorem, which indicates the prediction error of the many-model ensemble should be less than the average error of the individual models due to the diversity of predictions among the individual models. To investigate the implications of the No Free Lunch and Diversity Prediction Theorems, we developed a naïve ensemble-average model, which equally weights the predicted phenotypes of individual models. We evaluated this model using two traits influencing crop yield—days to anthesis and tiller number per plant—in the Teosinte Nested Association Mapping dataset. The results show that the ensemble approach increased prediction accuracies and reduced prediction errors over individual genomic prediction models. The advantage of the ensemble was derived from the diverse predictions among the individual models, suggesting the ensemble captures a more comprehensive view of the genomic architecture of these complex traits. These results are in accordance with the expectations of the Diversity Prediction Theorem and suggest that ensemble approaches can enhance genomic prediction performance and accelerate genetic gain in crop breeding programs.

**Article summary:** This research targets selective breeding industries and researchers developing genomic prediction models to accelerate genetic gain in breeding programs. We applied the concept of an ensemble, combining multiple individual genomic prediction models, to predict key traits (days to anthesis and tiller number per plant) in a crop breeding dataset. Here, we show that an ensemble approach increased prediction accuracies and reduced prediction errors over individual genomic prediction models. These results indicate the potential for ensembles of multiple, diverse genomic prediction models to accelerate genetic gain in breeding programs by increasing the accuracy of selection decisions.

## Introduction

Genomic selection has accelerated the rates of genetic gain in plant breeding programs (Voss-Fels *et al*. 2019) by using prediction models to associate genetic markers with trait phenotypes (Meuwissen *et al*. 2001). The ability to predict and select plants based on genetic markers, instead of trait phenotypes, has enabled novel selection schemes and breeding program designs with the potential to accelerate rates of genetic gain (Heffner *et al*. 2009; Gaynor *et al*. 2017; Powell *et al*. 2020). As of today, genomic selection has accelerated the rates of genetic gain for grain yield in commercial breeding programs (Cooper *et al*. 2014) and the integration of genomic selection in public breeding programs is underway (Dreisigacker *et al*. 2021; Prasanna *et al*. 2021).

The accuracy of genomic selection is dependent on the choice of a genomic prediction model. Therefore, research evaluating alternative genomic prediction models has received considerable attention. The primary goal of these research investigations is to develop a genomic prediction model that can consistently reach higher prediction performance compared with others. A major challenge in enhancing these models is capturing complex genetic interactions, often resulting from gene regulatory networks (Mascher *et al*. 2024). Cooper *et al*. (2005) demonstrated that epistatic (nonlinear) marker interactions can decrease the rate of genetic gain compared to the scenarios where only additive (linear) effects define the genetic architecture of complex traits. This finding emphasises the necessity to capture complex genetic interactions to maximise rates of genetic gain. Mackay (2014) discussed the possibility of performance improvement in genomic prediction by including epistatic effects in prediction models. Previous research (Montesinos-López *et al*. 2018; Pérez-Enciso and Zingaretti 2019; Washburn *et al*. 2021) has highlighted the difficulty of finding aa single genomic prediction model that sufficiently captures complex interactions to consistently outperform other models.

Outside of plant breeding, the absence of a best prediction model has been explained by the No Free Lunch Theorem (Wolpert and Macready 1997). The No Free Lunch Theorem postulates that the average prediction performance of individual models becomes equal over prediction problem scenarios. If we accept that this theorem applies generally, seeking superior individual genomic prediction models for the multiple prediction problems of plant breeding programs is unlikely to be successful. An alternative approach could be to generate ensemble combinations of multiple, different genomic prediction models.

Ensemble approaches combine predictions from multiple models (Page 2018; Farooq *et al*. 2022). Several ensemble approaches have been proposed, such as Bootstrap aggregating, which use multiple weak prediction models in parallel, such as Bootstrap aggregating (Breiman 1996), or sequentially, such as AdaBoost (Freund and Schapire 1995). These ideas informed the development of methods such as random forest (Breiman 2001a) and ensemble neural networks (Zhou *et al*. 2002; Li *et al*. 2004; Liu and Li 2008). Combining multiple distinctive prediction models is expected to cancel out errors derived from the gap between observed and actual values, leading to performance improvement (Page 2018; Kick and Washburn 2023).

Page (2018) provides a theoretical framework, illustrated with examples, of the potential advantages of applying ensembles of multiple, diverse models to enhance the prediction of key properties for complex multi-dimensional systems. One aspect of the framework is the “Diversity Prediction Theorem” which states that the model-ensemble error is equivalent to the average model error of the individual models minus the diversity of the model predictions. This theorem indicates prediction errors can be reduced, and prediction performance improved, whenever a suitable ensemble of diverse models can be identified. We consider the implications of the Diversity Prediction Theorem for applications of model ensembles for genomic prediction in crop breeding.

In general, ensemble approaches have been successfully leveraged in agricultural science. For instance, Wallach *et al*. (2018) demonstrated higher prediction accuracies of ensemble approaches for forecasting climate change scenarios in crop yield prediction. Other studies have shown the superiority of ensemble approaches over individual genomic prediction models (Bian and Holland 2015; McCormick *et al*. 2021; Huang and Wei 2022; Fradgley *et al*. 2023; Heilmann *et al*. 2023; Kick and Washburn 2023; Washburn *et al*. 2024). However, key factors leading to the performance increases of ensemble approaches for genomic selection have not been thoroughly investigated and discussed. Therefore, we investigate the performance of ensemble approaches for genomic selection in comparison to that of the individual genomic prediction models and dissect the factors leading to performance increases.

Here, we (1) investigate the prediction performance of six individual genomic prediction models for two traits controlled by a genetic architecture involving both linear and nonlinear interactions, (2) consider the implications of the No Free Lunch Theorem and evaluate an ensemble approach to assess its effectiveness compared to individual genomic prediction models and (3) use the “Diversity Prediction Theorem” as a framework to investigate how the diversity of model predictions contributes to the prediction performance of ensembles.

## Materials and methods

### Dataset

The TeoNAM dataset (Chen *et al*. 2019) consists of five recombinant inbred line (RIL) populations derived from five backcross hybrid crosses of the maize (*Z. Mays*) line W22 and five teosinte inbred lines; wild teosinte types TIL01, TIL03, TIL11 and TIL14 from *Z. mays ssp. parviglumis* and TIL25 from *Z*.*mays ssp. mexicana*. The maize inbred line W22 was used as the female plant for crossing. Following the F1 cross, the F1 was backcrossed once to W22. Each cross generated one RIL population and thus five different RIL populations were generated; W22TIL01, W22TIL03, W22TIL11, W22TIL14 and W22TIL25. After the backcross, each RIL population was advanced four times by controlled self-pollination to produce the RILs used for the analyses. Each RIL population was measured for seven agronomic traits and fifteen domestication traits. From the full trait list, days to anthesis (DTA) and tiller number per plant (TILN) were chosen as target traits for the ensemble investigations as both traits are expected to be a consequence of genetic interactions in a biologically complex network (Dong *et al*. 2012; DeWitt *et al*. 2021; Powell *et al*. 2022).

The traits were measured in two different environments. For W22TIL01, W22TIL03 and W22TIL11, the experiment was conducted in 2015 and 2016 summer. W22TIL14 was grown in 2016 and 2017 summer and W22TIL25 was evaluated in two blocks in 2017 summer. All the experiments were conducted at the University of Wisconsin West Madison Agricultural Research Station with a randomized complete block design.

The summary of genetic marker (SNP) and RIL numbers per cross is described in Table 1. Each RIL population contains at least 200 RILs with more than 10,000 SNPs.

**Table 1.** The number of SNPs in each preprocessing phase and recombinant inbred lines (RILs) in each population.

| Population<br>Name | SNPs |  |  | RIL |
| --- | --- | --- | --- | --- |
|  | Original | Imputation | LD Filtering | Number |
| W22TIL01 | 13,089 | 13,042 | 274 – 322 | 222 |
| W22TIL03 | 16,110 | 16,076 | 295 – 342 | 270 |
| W22TIL11 | 13,188 | 13,153 | 268 – 314 | 219 |
| W22TIL14 | 11,396 | 11,375 | 270 – 320 | 230 |
| W22TIL25 | 14,885 | 14,857 | 294 - 341 | 308 |

### Data cleaning & preprocessing

Preliminary quality control on the TeoNAM dataset identified three potential issues that required further investigation prior to the application of prediction models; (1) missing genomic marker calls, (2) missing trait measurements/records and (3) a larger number of genomic markers compared to the number of RILs.

Missing marker calls was one noticeable problem affecting the quality of the TeoNAM dataset for evaluating ensemble prediction methodology. Imputation of missing marker calls was undertaken when possible using two different methods. Our objective was not to evaluate the merits of alternative imputation methods, rather we were interested in assessing the implications of any imputation approach on the outcomes of the ensemble prediction methodology in relation to the expectations of the Diversity Prediction Theorem (Page 2018). Therefore, we examined the impact of alternative imputation methods on the prediction diversity among individual prediction models and the consequences of changes in the prediction diversity for the performance of the ensemble of models. Herein we present the results based on one imputation method and present the comparable results for an alternative imputation method in the Supplementary Materials for purposes of comparison. The imputation of missing marker calls with the most frequent allele was the primary approach leveraged to output prediction results. For the alternative imputation approach (Supplementary Material), missing marker calls were imputed with flanking markers. If flanking markers on both sides possessed the same parental allele, markers with missing calls were imputed with the same parental allele of their flanking markers. If flanking markers on both sides contained different parental alleles, the parental allele of the closest flanking marker was leveraged to impute the missing marker calls. If the alleles of markers in a chromosome were missing entirely, corresponding RILs were removed. For both imputation approaches, SNPs with more than 10% missing marker calls were removed.

A missing target trait problem occurred when the phenotype of a target trait was missing. Imputation was an infeasible choice in this dataset due to a lack of external information enabling the reasonable imputation of missing phenotypes. Hence, RILs without target trait phenotypes were excluded in this study.

The high proportion of markers relative to the number of RILs in each cross could negatively affect the performance of some of the genomic prediction models. The number of markers in all the crosses was considerably higher than the number of RILs (Table 1). The number of RILs may not have been sufficient to capture some of the complex patterns among markers and phenotypes, resulting in the curse of dimensionality (Bellman 1957; Ramstein *et al*. 2019). This can have more influence on the prediction performance of the machine learning models. Secondly, genetic markers may not provide additional information to increase prediction performance if they are in strong linkage disequilibrium (LD) and are not independent of each other. Such genetic markers can be removed to exclude redundant information. Additionally, training machine learning models with data containing many attributes can increase computational time, which can be prohibitive for complex machine learning models such as the graph neural network (GAT) used in this study. Therefore, to reduce the computation time for the machine learning models we reduced marker dimensionality by eliminating less informative markers based on their LD relationships. PLINK (v1.9) (Chang *et al*. 2015) was used to remove SNPs with a squared correlation of >0.8 using a window size of 30,000 bp and step size of 5. A higher correlation indicated that two genetic markers provided similar information for prediction. Thus, removing either of the genetic markers could be beneficial by reducing the total number of genetic markers rather than losing critical information for better predictions.

Since each RIL population was grown and measured in two distinctive environments, the sub-datasets were concatenated into a single dataset with a factor with two levels representing the different environments. This concatenated single dataset was randomly split into training and test sets for training and evaluating genomic prediction models, respectively. Genotype-by-environment (GxE) interactions were a source of uncertainty accounted for in the analyses. Investigations of the impact of GxE interactions in the genomic prediction models will be an area of focus for future research investigation.

The datasets after undertaking all the preprocessing steps were used as input for the six individual genomic prediction models. The final genetic marker (SNP) and RIL numbers used in this study are summarized in Table 1. The final number of SNPs varied depending on the combination of RILs in the training set for each sample.

### Diversity Prediction Theorem Framework

The effect of prediction model diversity on the prediction performance of an ensemble can be formulated in terms of the Diversity Prediction Theorem, as given by Page (2018):

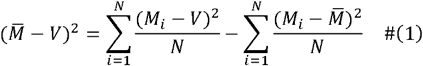

where *M*_*i*_ is the set of predicted values from prediction model *i*, 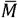 is the set of mean predicted values from the *i* individual prediction models, *V* the set of true values and *N* is the total number of prediction models considered. The Diversity rediction Theorem equation indicates that the many-is model error (the first term) equals the average error (second term) minus the prediction diversity (third term). Following Equation (1), the prediction diversity must be positive if the predictions differ. Hence, the many-model error must be smaller than the average error. Further, it is noted that as the prediction diversity decreases the third term approaches zero and the many-model error will become the average error.

In this study, we use the Diversity Prediction Theorem as a framework to investigate the potential of an ensemble of multiple genomic prediction models to enhance prediction performance in an empirical crop breeding dataset. Therefore, a few alterations in definitions were required to apply the Diversity Prediction Theorem to an empirical crop breeding dataset, such as the TeoNAM dataset. In our study, *M*_*i*_ was defined as predicted phenotypes from individual genomic prediction models. While *V* was defined as trait observations, instead of true values as per the original theorem. We refer to the many-model error as the ensemble error throughout this manuscript. Six individual genomic prediction models were applied in our analysis. Therefore, *N* = 6. We report the mean values, by trait, for the ensemble error (the first term), the average error (the second term) and the prediction diversity (the third term) across all prediction scenarios and by training-test set ratio.

### Individual genomic prediction models

Six prediction models, three classical genomic prediction and three machine learning models, were applied to the TeoNAM dataset. The three classical genomic prediction models were Ridge Regression Best Linear Unbiased Prediction (rrBLUP) (Meuwissen *et al*. 2001), BayesB (Meuwissen *et al*. 2001) and Reproducing Kernel Hilbert Space (RKHS) (Gianola and van Kaam 2008). The three machine learning models were Random Forest (RF) (Breiman 2001a), Support Vector Regression (SVR) (Drucker *et al*. 1996) and Graph Attention Network (GAT) (Brody *et al*. 2021).

### Classical genomic prediction models

#### Parametric models

Parametric models have been widely leveraged for genomic prediction (Meuwissen *et al*. 2001). One major characteristic in parametric models is the requirement of assumptions in the input distribution. Such assumptions allow the model parameters to be determined within a finite value range.

The most well-known parametric models in genomic prediction are linear mixed models. They have been leveraged since the initial developmental stage in genomic prediction (Ray *et al*. 2022). A linear mixed model, in general, can be formulated as Equation (2) (Pérez and de Los Campos 2014):

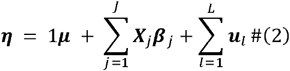

where ***η***= {*η*_1_, *η*_1_,…, *η*_*n*_} is a set of the predicted phenotypes for true values ***Y***= {*Y*_1_, *Y*_1_,…, *Y*_*n*_}, ***µ*** is intercept, ***X***_*j*_ is design matrices representing marker values, ***β***_*j*_ is coefficient matrices indicating each genomic marker effects and ***u***_***l***_ = { *μ* _*l* 1_, *μ*_*l* 1_,…, *μ*_*ln*_}is random effects in a vector format. The assumed value distribution for ***β***_*j*_ differs among models. The linear mixed models aim to determine ***β***_*j*_ in a way that the gap between ***η*** and ***y*** is minimum.

rrBLUP is a commonly used mixed linear model in genomic prediction, assuming that the effect of each marker is small and normally distributed). with the same variance regardless of the effect size of each genetic marker (Meuwissen *et al*. 2001; Bernardo and Yu 2007). Hence, ***β***_*j*_ is distributed as 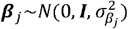 where ***I*** is an identity matrix and *σ*^2^ is the variance of genomic marker effects (Endelman 2011; Endelman and Jannink 2012). Genomic best linear unbiased prediction (GBLUP) (VanRaden 2008) is another linear mixed model emphasising the relationship between individuals for predictions by developing the genomic relationship (kinship) matrix (Lipka *et al*. 2012; Wang *et al*. 2015). This method is demonstrated to be equivalent to rrBLUP and both models share the same mechanisms and assumptions (Habier *et al*. 2007). Hence, only the rrBLUP model implementation was chosen as one of the genomic prediction models for the ensemble investigations in this study.

Another group of linear mixed models used for genomic prediction analyses is the Bayesian methods often characterized by the Bayesian alphabet series (Gianola *et al*. 2009). In contrast to rrBLUP that assumes all the markers have normally distributed small effects, Bayesian methods allow each genomic marker effect ***β***_*j*_ to have different distributions (Wang *et al*. 2018). This variation in the distribution is expected to capture more realistic genomic marker effects whenever it is unrealistic for all the genomic marker effects to have the same distribution with small effects, as is assumed in rrBLUP. Another noticeable characteristic of the Bayesian models is the application of prior distributions, constructed from prior knowledge, assumptions and statistics, iteratively updated by the accumulation of information from each sampling to capture the properties of the true distribution (Kruschke 2010). Among various Bayesian models, BayesB was selected in this study because it has been widely used as one of the standard models in genomic prediction (Abdollahi-Arpanahi *et al*. 2020; Farooq *et al*. 2022; John *et al*. 2022; Meher *et al*. 2022; Plavšin *et al*. 2022). For BayesB some genomic marker effects are expected not to be influential on target phenotypes and their genomic marker effects are set as zero.

We developed rrBLUP and BayesB models using the library BGLR (Pérez and de Los Campos 2014) in R. For parameter setting, we set the number of iterations and burn-in as 12,000 and 2,000 respectively for both models. Other parameters were set as default throughout this study.

#### Semiparametric models

Semiparametric models possess both parametric and nonparametric properties, indicating that the models can capture both linear and nonlinear relationships from two different approaches. One of the semiparametric models that has been widely leveraged for genomic prediction is a RKHS regression model introduced by Gianola and van Kaam (2008). The original idea of RKHS was proposed by Aronszajn (1950), which mapped given data into Hilbert space to capture complex nonlinear relationships among the data points. Any relationship that may not be captured in the original dimension can be observed on complex hyperplanes. Gianola and van Kaam (2008) leveraged RKHS to detect nonlinear effects represented as dominance and epistatic genetic effects, while maintaining the parametric components to capture the additive genetic effects. RKHS can be formulated as Howard *et al*. (2014) denoted with the base of Equation (2):

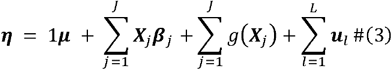

where *g*(***X***_*j*_) represents genetic effects from nonlinear genomic marker effects such as dominance and epistatic effects. *g*(***X***_*j*_) is expressed as

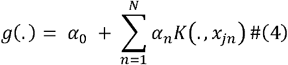

where ***X****j* = {*x*_*j* 1_, *x*_*j* 1_,…, *x*_*jn*_}, *α*_0_ is a fixed term, *x*_*n*_ is coefficient accompanied to *x*_*n*_ and *K* is the reproducing kernel. These equations imply that the RKHS model consists of both additive and non-additive components, aimed at capturing genomic marker effects in a more comprehensive way. Similarly, the BGLR library was leveraged for the RKHS model. A Gaussian kernel was used with a fixed bandwidth parameter. The number of iterations and burn-in are also set as 12,000 and 2,000 and the rest of the parameters remained as default as well.

#### Machine learning models

Machine learning (nonparametric) models do not require any assumptions about the underlying distributions of the model terms. Instead, the parameters of the models are determined by an iterative training process. From the various machine learning methods, we selected Random Forest (Breiman 2001a), Support Vector Regression (Drucker *et al*. 1996) and Graph Attention Network (Velickovic *et al*. 2017) for our investigation of ensemble prediction.

RF contains a collection of decision trees (Belson 1959) trained by sub-train sets sampled from the original training set (Liu *et al*. 2012). Decision trees develop a tree-like decision mechanism flow, consisting of nodes and edges. After two edges are released from the top node called the root, each layer consists of nodes with one incoming and two outgoing edges except the end layer nodes called leaves which have no outgoing edges (Rokach and Maimon 2005). Each node except the leaves holds a condition based on values in a specific feature from the given data. The algorithm starts traversing from the root, and if the target data point (a set of features) satisfies the condition of the root, the algorithm traverses the tree to the left bottom adjacent node and the right bottom adjacent node in vice-versa. This process repeats until it reaches the leaves determining the class or value of the target data point (de Ville 2013). When a target data point is given, each decision tree returns a prediction value, and the final prediction value is determined by aggregating the prediction results from all the trees. Using different sub-train sets for training, respective decision trees can cancel out prediction noise from each tree, resulting in more stable prediction results (Qi 2012). RF was chosen as a model because it is one of the commonly used machine learning models in genomic prediction (González-Camacho *et al*. 2018; Sandhu *et al*. 2021a; Farooq *et al*. 2022; John *et al*. 2022).

SVR is another machine learning approach that draws a hyperplane between data points for continuous target values. A hyperplane is drawn in a way that the distance between the hyperplane and the closest data points (support vector) becomes maximum with minimum prediction errors that include the largest number of data points within the range of the decision boundary. This is formalized by minimising the following objective function under several constraints (Drucker *et al*. 1996):

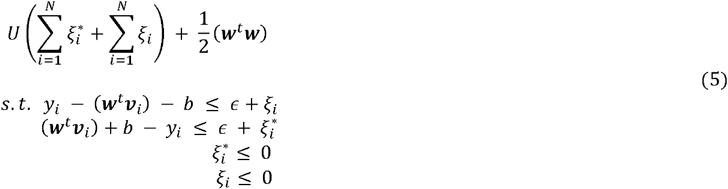

where *U* is an objective function targeted for the constraints, *b* is an intercept, ***w*** is a coefficient vector for a feature vector, *ϵ* is the distance between a hyperplane and a decision boundary, *υ* is a vector of data points, 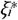 and ξ_*i*_ are slack variables functioned to make the decision boundary “soft” by allowing some data points to be outside the upper and lower boundary. SVR can be equipped with a kernel, mapping data points to another dimension for capturing nonlinear interactions. SVR has also been leveraged for various experiments in genomic prediction as a standard method (An *et al*. 2021; Yu *et al*. 2021; John *et al*. 2022; Li *et al*. 2023) and is selected as one prediction model in this study.

GAT is a graph neural network that applies a self-attention mechanism for predictions. We apply GAT by converting the relationship between markers and phenotypes into a graphical format. Genetic markers and phenotypes can be represented as nodes, and the connections between genetic marker nodes and phenotype nodes are represented as edges. The edges are directed from the genetic marker nodes to phenotype nodes, showing that genetic markers explicitly affect phenotypes. In this study, we do not add connections between marker nodes because allowing the edges between them did not improve the prediction performance and resulted in the exponential increase of computational time. We leverage the GAT model proposed by Brody *et al*. (2021). The attention mechanism can be written as below:

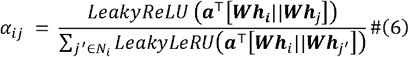

where ***a***^T^∈ *R*^2*d*′^ is a transposed weight vector, *d*′ is the number of features in each node,***W*** is a weight matrix, *h*_*j*_ **=**{ *h*_1_, *h*_*2*_,…, *h*_*N*_}is a set of node features of node *i*, ‖ concatenates vectors, *j*∈ *N*_*i*_ is a partial neighbor of nodes sampled from the entire neighbors. A Leaky Rectified Linear Unit (LeakyReLU) activation function was applied to convert calculated attention values nonlinearly. Unlike Rectified Linear Unit (ReLU) which returns zero for values smaller than zero, this multiplies the attention values with a slope (0.01 in this study) if the given values are below zero. The feature values of neighbors and trainable weights are nonlinearly activated to generate the attention values, and the calculated attention value is normalized at the end. This attention calculation mechanism is repeated for *K* times to stabilise the calculation result. It is called multi-head attention and is formulated as below (Velickovic *et al*. 2017):

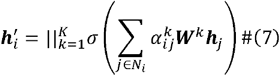

where *K* is the number of total attention layers, *σ* is a nonlinear activate function and 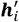 is the updated node features. At the final layer, the result from each attention layer is averaged instead of concatenation:

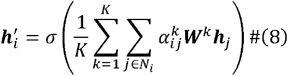

This final value 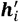 is used as a predicted phenotype from the model. GAT was chosen among graph neural networks due to its attention mechanism which can more precisely capture key prediction patterns underlying the given data to enhance prediction performance.

For RF and SVR, Sklearn (v1.2.2) was used for the model implementation in Python. The number of trees in RF was set as 1,000 while the default setting was used for other hyperparameters. In SVR, the radial basis function (RBF) was used for the kernel and the default setting was used for the remaining parameters. For GAT, Pytorch Geometric (v2.3.0) was used. Since phenotype and genetic marker nodes contained different types of information, they needed to be identified as different node types. Hence, the graph was converted into a heterogeneous graph. The model was three-layered with one hidden layer with 20 channels and a dropout rate of 0 was applied to every layer. The Exponential Linear Unit (ELU) function was used for the activation functions. The total number of heads was set as 1. The model was trained with 50 epochs by mini-batched graphs with a batch size of 8. AdamW was selected for the optimiser with a learning rate of 0.005 and weight decay of 0.

### Naïve ensemble-average model

The naïve ensemble-average model leveraged in this study is formulated as below:

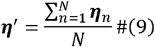

where 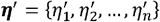 is a vector of the final predicted phenotypes, ***η*** represents predicted phenotypes from each individual genomic prediction model and *N* is the total number of individual genomic prediction models, as defined in equation (1), that was assigned as 6 in this study. Equation (9) indicates predicted phenotypes from each individual genomic prediction model were averaged with the same weight.

### Genomic marker effect estimation

The extraction of estimated genomic marker effect values from each model can suggest how each model estimates the genomic marker effect of respective SNPs for predicting target phenotypes, allowing a comparison of the genomic prediction models at the genomic level. For rrBLUP and BayesB, the genomic marker effects were extracted using ***β*** which represents allele substitution effect.

For RKHS and SVR, the Shapley value (Shapley *et al*. 1953) was employed to estimate genomic marker effects. The Shapley value is a metric, originally in the field of game theory, to assess the equitable distribution of resources and rewards to players based on their contribution level under cooperative scenarios (Winter 2002). A larger Shapley value is allocated to SNPs causing larger changes in predicted values. The SNP Shapley value is calculated below (Lundberg and Lee 2017):

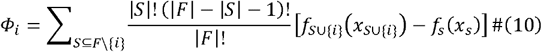

where *Φ*_*i*_ is a Shapley value for feature *i*(*SNP*_*i*_ in this case), *S* is a subset of features *F, f*_*SU*{*i*}_is a model predicting a target value with a feature subset including *i, f*_s_ is a model predicting the target value without including. In this study, the Shapley value, a genomic marker effect of a SNP, is estimated as the average gap in predicted phenotypes between the cases where the target SNP is included and the case where the target SNP is excluded. For RKHS, the Shapley value was implemented using iml (v0.11.3) (Molnar *et al*. 2020) in R whereas SHAP (v0.42.1) (Lundberg and Lee 2017) was applied to SVR in Python. Since the Shapley value returns element-wise feature effects, the final Shapley value of each SNP is calculated by averaging the Shapley values for the target SNP from RILs in a test set.

For RF, genomic marker effects were estimated by extracting impurity-based feature importance values, measuring the importance of features by splitting the data using a value *s* from *X* at node *t* formulated as below (Ishwaran 2015):

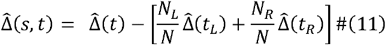

where 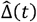 represents impurity at node *t, N* is the total number of data points in daughter node *t*_*L*_ and *t*_*R*_, 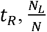 is the weight of left daughter node *t*_*L*_, 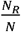 is the weight of right daughter node *t*_*R*_, 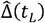is impurity value of left daughter node *t*_*L*_ and 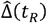 is impurity value of right daughter node. The impurity is measured by the summed squared value at the gap between the mean value of the prediction target and the actual value of each element in each node. Since this denotes only the impurity gap at node *t*, the summation of the impurity gap across all nodes leads to the total impurity gap for a target feature SNP. The importance of the feature was extracted using Sklearn (v1.2.2) in Python.

The genomic marker effects of GAT were estimated by leveraging an interpretability method called Integrated Gradients (Sundararajan *et al*. 2017), integrating the gradient of a line drawn between two points; the baseline point shows the prediction value without the effect of the target feature SNP and the other indicates the prediction value with the feature SNP value. Using the baseline point, the true gradient can be calculated by eliminating the effect of the initial value. Integrated Gradients is calculated as below (Sundararajan *et al*. 2017):

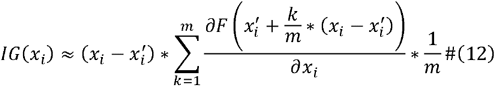

where *x* is a feature, *x*^′^ is a baseline value of *x* and *m* is the total number of interpolation steps of the integral. Similar to the Shapley value, Integrated Gradients also returns element-wise marker effect, and thus the final genomic marker effect was estimated by averaging the effect of the target SNP across RILs in a test set. Pyg (v2.4.0) was used for the implementation.

## Assessment criteria

### Experimental flow

The experimental framework for evaluation of the individual genomic prediction and the ensemble models is shown in Figure 1. This study evaluated the performance of the genomic prediction models under within-population prediction scenarios. Each RIL population, data in the TeoNAM dataset was randomly split into training and test sets. After SNPs were filtered based on the LD filtering, the training set was used to train the six individual genomic prediction models. The performance of the trained individual genomic prediction models was evaluated using the test set. Vectors of predicted phenotypes for RILs in the test set, derived from each individual genomic prediction model, were assembled to construct a predicted phenotype matrix. The ensemble genomic prediction model leveraged the predicted phenotype matrix as the input. After evaluating the prediction performance of the ensemble approach, the genomic marker effects from each of the six individual genomic prediction models were extracted by the interpretable approaches explained in Section Genomic marker effect estimation.

**Figure 1.**
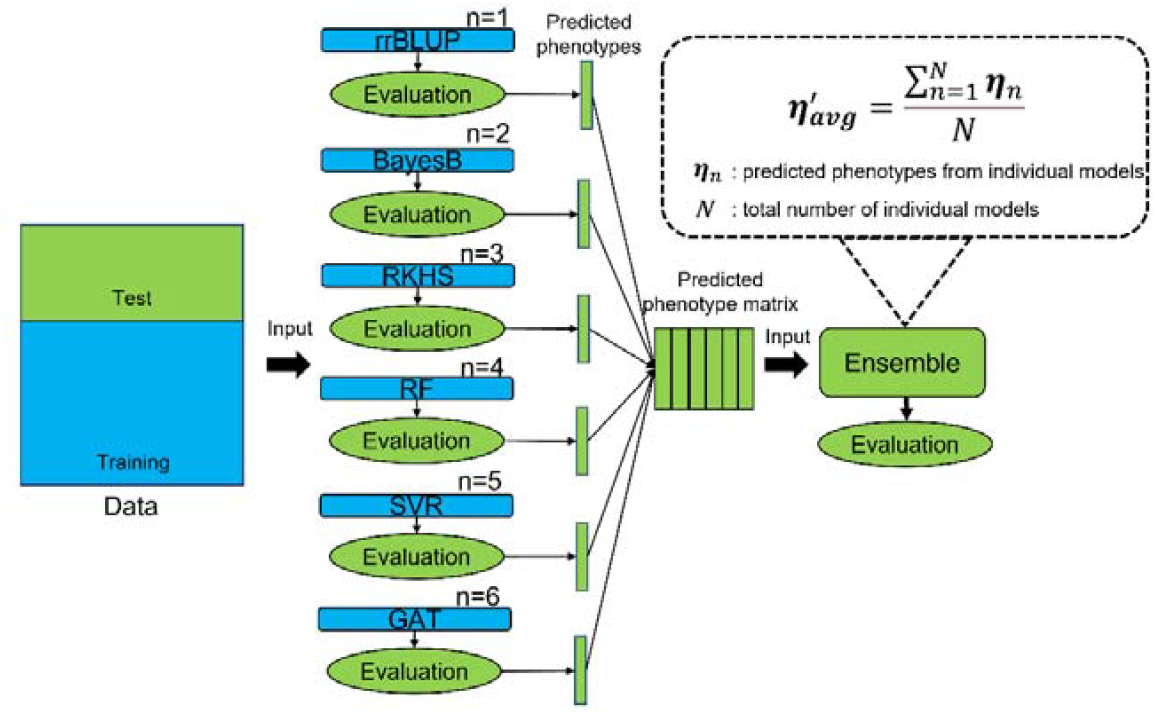
A diagram of the experimental flow used in this study. The blue and green colors represent components related to a training and test set, respectively. Each individual genomic prediction model was trained using genetic markers. The performance of the trained individual genomic prediction models was evaluated using the test set. Predicted phenotypes for RILs in the test set from the individual genomic prediction models were used as input data for the ensemble model.

To ensure the generalisability of the observed result, we repeatedly implemented the genomic prediction models for several different settings. Three different training-test ratios (0.8-0.2, 0.65-0.35 and 0.5-0.5) were leveraged with a random sampling of 500. Hence, each genomic prediction model was evaluated over 1,500 prediction results (3 ratios*500 samples) in each population-trait combination.

### Metrics

Two metrics were leveraged to measure the performance of the genomic prediction models. The Pearson correlation measured the concordance of ranks between the predicted and observed phenotypes for the test set to measure prediction accuracy (1 indicates that the ranking between predicted and observed phenotypes completely matches). Mean squared error (MSE) was used to measure the prediction error between the predicted and observed phenotypes (0 indicates that there is no difference between the predicted and observed phenotypes).

## Results

### Lower ensemble prediction error than average model prediction error with diverse models

An improvement in prediction performance was observed using the naïve ensemble-average model compared to the average of individual genomic prediction models. As shown in Figure 2, The naïve ensemble-average model outperformed the average of the individual genomic prediction models in prediction accuracy (Pearson correlation) and mean squared error (MSE) for both DTA and TILN traits. The median prediction accuracy of the naïve ensemble-average model (0.919 for DTA and 0.790 for TILN) was higher than the average of the individual genomic prediction models (0.719 for DTA and 0.671 for TILN). For prediction error, the naïve ensemble-average model reached a lower median MSE (10.167 for DTA and 0.277 for TILN) compared to the average of the individual genomic prediction models (16.893 for DTA and 0.356 for TILN).

**Figure 2.**
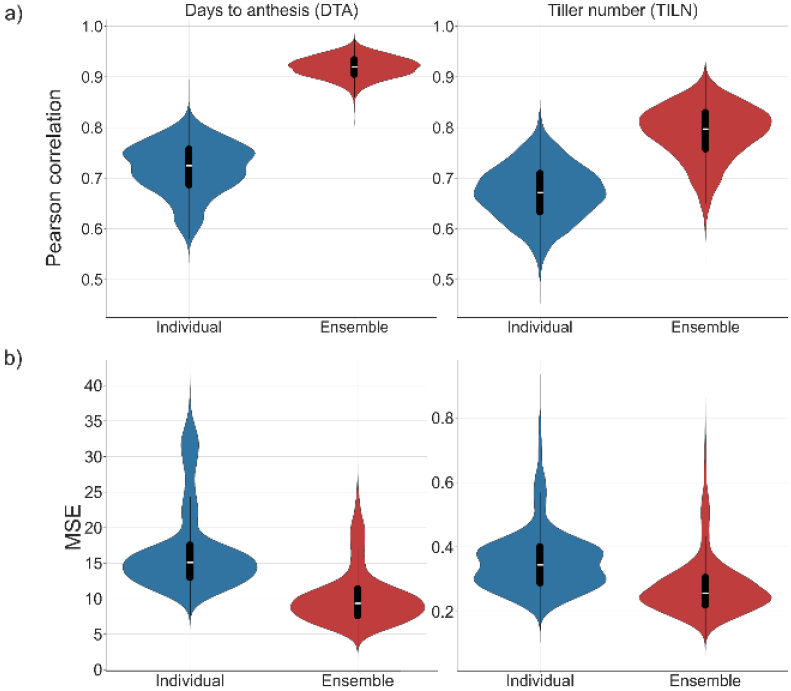
Violin plots comparing genomic prediction performance of the average of individual genomic prediction models (Individual) in the blue versus the naïve ensemble-average model (Ensemble) in the red. The performance of genomic prediction models was measured with a) the Pearson correlation and b) mean squared error (MSE). The width of the violins represents the distribution of metric values for predictions from all combinations of the five RIL populations, three traini g-test ratios and 500 random samples. Box plots within the violin plots represent the median metric value (white line) and the interquartile range (black box) with whiskers extending 1.5 times the interquartile range.

Figure 3 visually represents the diversity in prediction performance among each individual genomic prediction model. For prediction accuracy, the median Pearson correlation of rrBLUP, BayesB, RKHS, RF, SVR and GAT was 0.878, 0.887, 0.767, 0.782, 0.312 and 0.802 for the DTA trait and 0.705, 0.708, 0.657, 0.668, 0.605 and 0.665 for the TILN trait. For the prediction error, the median MSE of rrBLUP, BayesB, RKHS, RF, SVR and GAT was 8.715, 8.048, 17.217, 14.268, 35.613 and 17.500 for the DTA trait and 0.319 and 0.315, 0.365, 0.357, 0.401 and 0.392. This demonstrates prediction diversity among the individual genomic prediction models, creating the potential to reduce the ensemble error.

**Figure 3.**
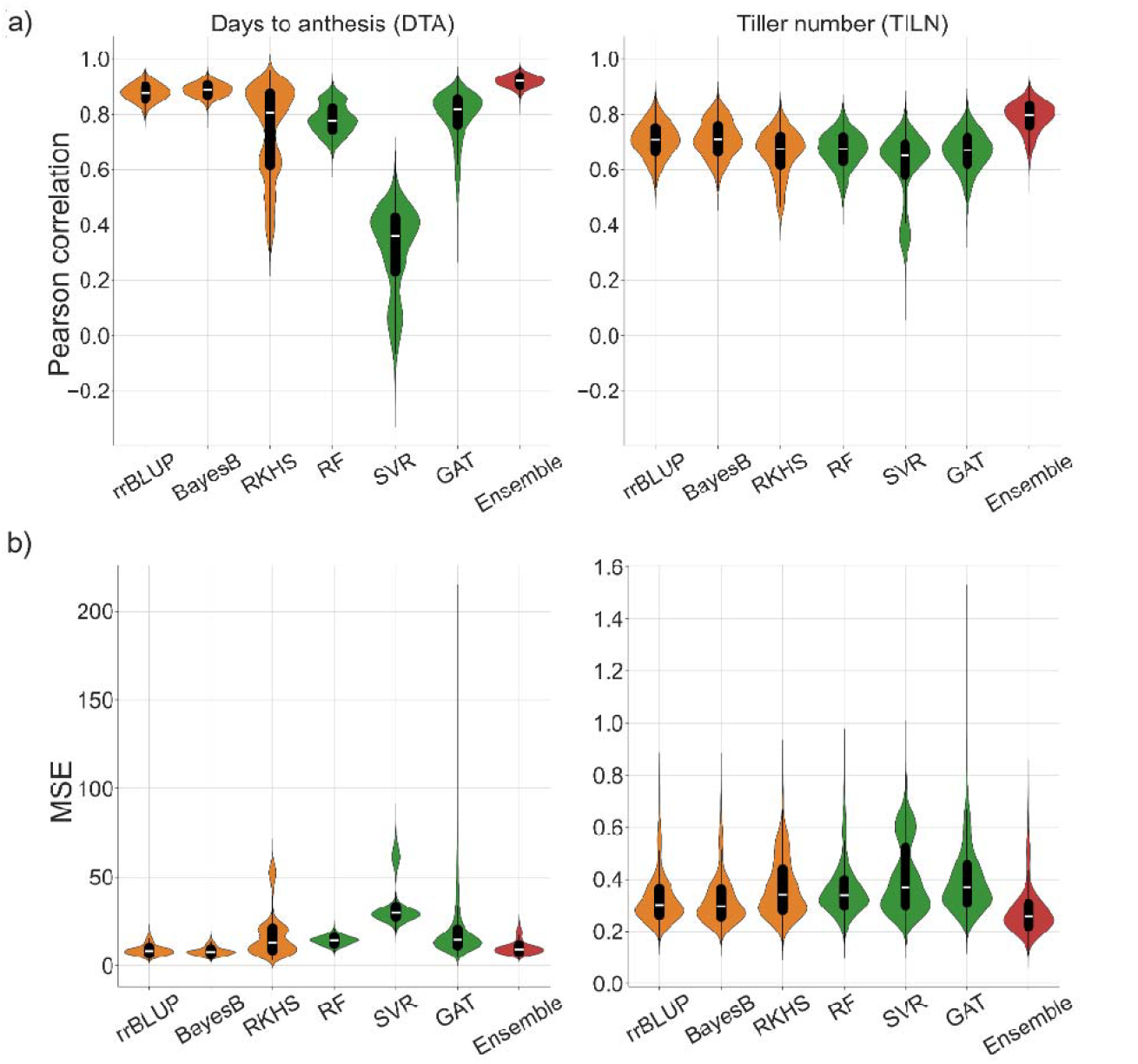
A comparison of genomic prediction performance of the naïve ensemble-average (Ensemble) model versus each of the individual genomic prediction models in violin plots. The width of the violins indicates the distribution of the metric values for predictions from all combinations of the five RIL populations, three training-test ratios and 500 random samples. The performance of genomic prediction models was measured with a) the Pearson correlation and b) mean squared error (MSE). The orange represents the performance of classical models (rrBLUP, BayesB and RKHS) while the green represents machine learning models (RF, SVR and GAT). The red is the performance of the ensemble. Box plots within the violin plots represent the median metric value (white line) and the interquartile range (black box) with whiskers extending 1.5 times the interquartile range.

The advantage of the naïve ensemble-average model over the individual models indicates that the prediction diversity among individual genomic prediction models was sufficient to decrease the ensemble error compared to the average error (Table 2). The ensemble error was lower than the average error for both traits. The value for the ensemble error was 10.17 for the DTA trait and 0.28 for the TILN trait, while the average error was 17.26 for DTA and 0.36 for TILN. The lower ensemble error was attributed to the prediction diversity among the individual genomic prediction models, which was 7.09 and 0.09 for the DTA and TILN traits, respectively (Table 2).

**Table 2.** Estimates of the average value and their standard error for each term of the Diversity Prediction Theorem (Equation (1)) across 7,500 scenarios for the traits days to anthesis (DTA) and tiller number per plant (TILN)

|  | ensemble error<br>(first term) | average error<br>(second term) | prediction diversity<br>(third term) |
| --- | --- | --- | --- |
| DTA | 10.17±1.34 | 17.26±1.77 | 7.09±0.73 |
| TILN | 0.28±0.05 | 0.36±0.05 | 0.09±0.01 |

Considered in terms of the Diversity Prediction Theorem, these results indicate that for both traits measured in the TeoNAM data set, the naïve ensemble-average model improved prediction performance by reducing the ensemble error compared to the average error with diverse individual genomic prediction models (Figures 2, 3, Table 2).

### Ensemble of models outperformed the best individual genomic prediction models

The naïve ensemble-average model demonstrated higher prediction accuracies and lower prediction errors compared to the best individual genomic prediction models for both traits when averaged across populations and training-test ratios (Figure 3).

For the DTA trait, the best genomic prediction model depended on which metric (prediction accuracy or prediction error) was used for the performance evaluation. Prediction accuracy was highest for the naïve ensemble-average model (median = 0.920) with the best individual genomic prediction model being BayesB (median = 0.888). In contrast, BayesB reached the lowest prediction errors (median = 7.765) among all the genomic prediction models including the naïve ensemble-average model (median = 9.340).

For the TILN trait, the highest prediction accuracies and lowest prediction errors were observed with the naïve ensemble-average model. The highest prediction accuracy was observed in BayesB (median = 0.709) within the individual genomic prediction models and the naïve ensemble-average model surpassed it (median = 0.797). BayesB reached the lowest prediction error among the individual genomic prediction models (median = 0.297), but the lower prediction error was observed in the naïve ensemble-average model (median = 0.257). The same trend was observed at the per-population level (Figure S1) and per training-test ratio (Figure S2).

### No consistent winner amongst individual genomic prediction models

The individual genomic prediction model with the highest prediction accuracies and the lowest prediction errors varied across the three training-test ratios (Table 3). This result suggests the absence of a “best” individual genomic prediction model for all prediction scenarios.

**Table 3.** Percentage of best performance achieved by each genomic prediction model in the respective training-test ratio (0.8, 0.65, 0.5) and trait (DTA, TILN) combinations. The performance was measured by Pearson correlation and MSE. The value reaching the highest percentage in each combination is highlighted in bold.

| Ratio | Trait | Pearson correlation |  |  |  |  |  |  | MSE |  |  |  |  |  |  |
| --- | --- | --- | --- | --- | --- | --- | --- | --- | --- | --- | --- | --- | --- | --- | --- |
|  |  | rrBLUP | BayesB | RKHS | RF | SVR | GAT | Ensemble | rrBLUP | BayesB | RKHS | RF | SVR | GAT | Ensemble |
| 0.8 | DTA | 1.24 | 0.44 | 0.40 | 0.00 | 0.00 | 0.00 | **97.92** | **39.32** | 31.92 | 15.64 | 0.08 | 0.00 | 0.00 | 13.04 |
|  | TILN | 0.00 | 0.40 | 0.00 | 0.00 | 0.04 | 0.00 | **99.56** | 2.84 | 9.28 | 0.00 | 0.04 | 0.24 | 0.04 | **87.56** |
| 0.65 | DTA | 0.24 | 1.08 | 0.00 | 0.00 | 0.00 | 0.00 | **98.68** | 18.12 | **72.00** | 0.12 | 0.04 | 0.00 | 0.00 | 9.72 |
|  | TILN | 0.00 | 0.24 | 0.00 | 0.00 | 0.08 | 0.00 | **99.68** | 1.00 | 13.64 | 0.00 | 0.00 | 0.08 | 0.04 | **85.24** |
| 0.5 | DTA | 0.04 | 2.08 | 0.00 | 0.00 | 0.00 | 0.00 | **97.88** | 4.44 | **94.12** | 0.00 | 0.20 | 0.00 | 0.00 | 1.24 |
|  | TILN | 0.00 | 0.24 | 0.00 | 0.00 | 0.00 | 0.00 | **99.76** | 0.24 | 16.04 | 0.00 | 0.00 | 0.00 | 0.00 | **83.72** |

Among the individual genomic prediction models, BayesB maintained the highest prediction accuracy percentage across training-test sets for the three training-test ratios (DTA ranged between 0.44% and 2.08% and TILN between 0.24% and 0.40%). While the second highest prediction accuracy percentage in the individual genomic prediction models was rrBLUP with the range of 0.04% and 1.24% for the DTA trait and 0.00% for the TILN trait. The highest prediction accuracy percentage was observed with the other individual genomic prediction models (RKHS, RF, SVR and GAT) in less than 1.00% of training-test set samples.

The individual genomic prediction model with the highest percentage of lowest prediction errors depended on the combination of training-test ratios and traits. For the DTA trait, the lowest prediction error percentage was the highest in rrBLUP (39.32%) when the training-test set ratio was 0.8, but BayesB was the highest when the ratio of the training set became smaller (72.00% for 0.65 and 94.12% for 0.5). For the TILN trait, the lowest prediction errors among the individual genomic prediction models were observed with BayesB across all ratios of the training-test (from 9.28% to 16.04%).

### Classical genomic prediction models outperform machine learning models

Figure 3 indicates classical genomic prediction models (rrBLUP, BayesB and RKHS) demonstrated higher prediction accuracies and lower prediction errors than machine learning models (RF, SVR and GAT). However, the magnitudes of differences were trait-dependent.

For the DTA trait, SVR had considerably lower median Pearson correlations (0.360) and higher median MSE (29.811) than other individual genomic prediction models. RF and GAT demonstrated comparable prediction accuracies and errors to RKHS with median Pearson correlations (0.778, 0.818 and 0.806) and median prediction errors (14.274, 14.537 and 12.976). rrBLUP and BayesB demonstrated the highest median prediction accuracies (0.878 and 0.888) and lowest median prediction errors (8.308 and 7.765) of all the individual genomic prediction models.

For the TILN trait, smaller performance differences between the classical genomic prediction models and machine learning models were observed in both prediction accuracy and error. SVR demonstrated the lowest median Pearson correlations (0.650) and higher median MSE errors (0.368) compared to the other individual genomic prediction models. RF and GAT demonstrated comparable prediction accuracies and errors to RKHS with median Pearson correlations (0.676, 0.670 and 0.675) and median MSE errors (0.339, 0.369 and 0.340). rrBLUP and BayesB demonstrated the highest median prediction accuracies (0.708 and 0.709) and lowest prediction errors (0.302 and 0.297) of all the individual genomic prediction models.

### Large variation in the genomic marker effects estimated by individual genomic prediction models

Large magnitudes of variation were observed in genomic marker effects across the classical and machine learning genomic prediction models. Figure 4 shows the pairwise comparisons of predicted phenotypes (top right triangle) and genomic marker effects (bottom left triangle) between the individual genomic prediction models. Positive associations were observed between several pairwise comparisons of predicted phenotypes, but not consistently observed for comparisons of genomic marker effects. Only a few models showed positive associations were between genomic marker effects. Each individual model estimated large traits effects to different SNP markers, leading to diversity in the estimated genomic marker effect sizes. Despite the variation in genomic marker effect sizes, the SNPs identified as putative QTL by Chen *et al*. (2019) were frequently included as features in the individual genomic prediction models (Figure 4).

**Figure 4.**
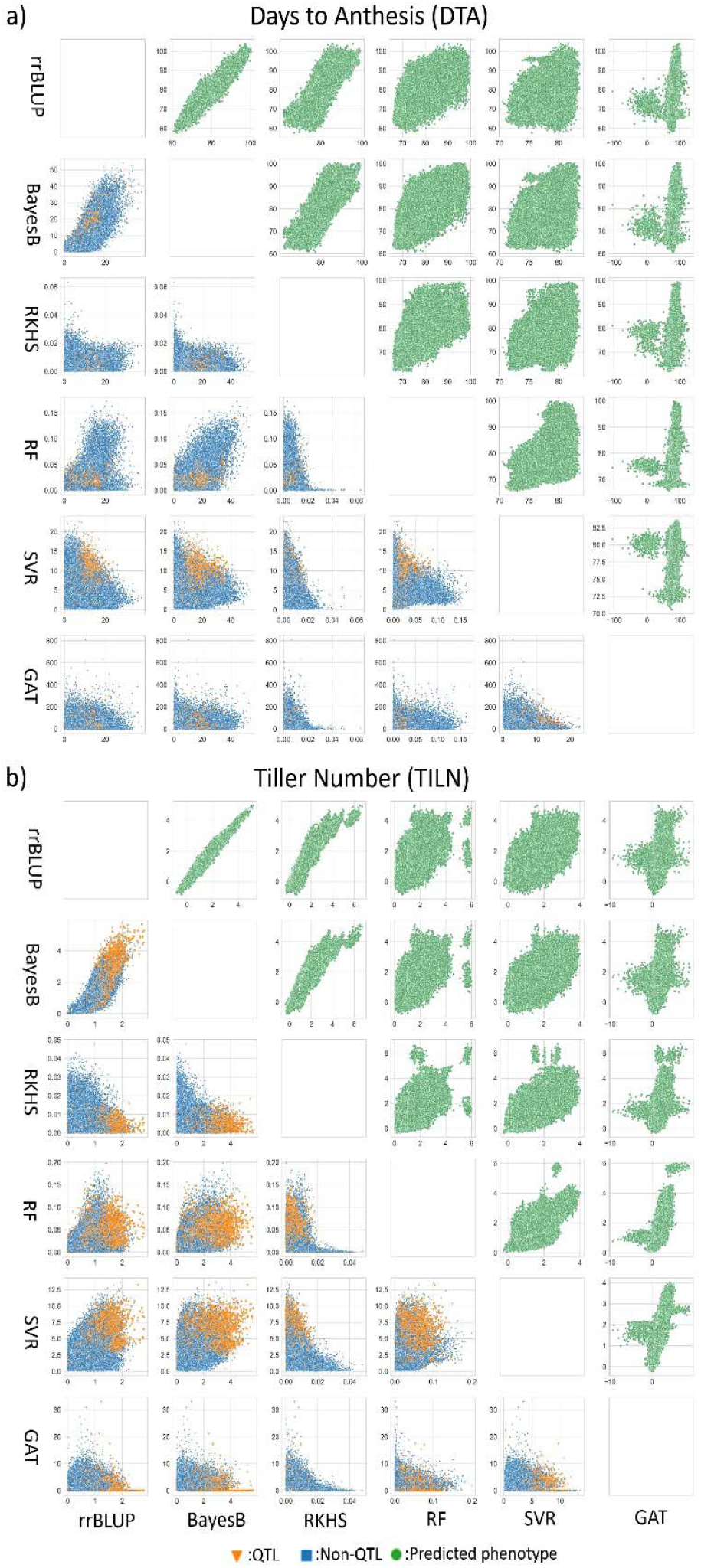
Pairwise comparison of individual genomic prediction models at predicted phenotypes (top right triangle) and genomic marker effects (the bottom left triangle) levels for both traits; a) the days to anthesis (DTA) and b) the tiller number per plant (TILN). The green circle dots represent a pair of predicted phenotypes for RILs included in the test set in each sample scenario. The blue square and orange triangle dots indicate a pair of estimated genomic marker effects in each sample scenario classified as non-QTL and QTL by Chen *et al*. (2019), respectively. A genetic marker was classified as a QTL if it was the closest to a QTL position within the support interval of 2 logarithm of the odds (LOD) calculated by Chen *et al*. (2019). Each point represents predicted phenotypes or genomic marker effect of each SNP in predictions from all combinations of the five RIL populations, three training-test ratios and 500 random samples.

For the predicted phenotypes, the strength of the associations among the individual genomic prediction models did not vary significantly (Table S1). Strong positive associations were observed among the classical genomic prediction models, with high Pearson correlations between rrBLUP and BayesB, rrBLUP and RKHS and BayesB and RKHS. Weaker but positive associations were observed among machine learning models, with high Pearson correlations between RF and SVR, RF and GAT and SVR and GAT. Furthermore, strong positive associations were observed among classical and machine learning genomic prediction models.

In contrast, lack of associations between genomic marker effects were consistently observed among individual genomic prediction models. While a relatively high association was observed between rrBLUP and BayesB, most other pairs showed weak associations, with Pearson correlations lower than 0.5 between the classical, machine learning and mixed genomic prediction models for both traits.

## Discussion

### Prediction performance depends on the complexity of the network affecting a trait

The complexity of an underlying biological network controlling a target trait can contribute to differences in prediction performance (Cooper *et al*. 2005). While networks of genes control the DTA trait (Buckler *et al*. 2009; Dong *et al*. 2012), models based on additive effects alone have been sufficient to account for the phenotypic diversity. In contrast, the TILN trait results from nonlinear marker interactions of the shoot branching network (Doebley *et al*. 1995; Bertheloot *et al*. 2020; Powell *et al*. 2022). The genomic prediction models evaluated in this study may lack mechanisms to fully capture patterns of genetic variation generated by such intricate networks. Azodi *et al*. (2019) compared the prediction performance of parametric (rrBLUP and Bayesian) against nonparametric machine learning models (RF, SVR and neural networks) for complex traits such as crop yield and plant height across various crops. Across the genomic prediction models evaluated, low prediction accuracies were consistently observed for many traits, indicating that the high complexity of gene networks underlying target traits in crop breeding can reduce the predictive performance of genomic prediction models.

The difference in the complexity of networks underlying target traits can also inferred by comparing the performance of individual genomic prediction models. For the DTA trait, nonparametric machine learning models (RF, SVR and GAT) showed lower prediction performance than the parametric models (rrBLUP and BayesB), especially for SVR. However, this lower prediction performance diminished for the TILN trait. The different prediction performances of parametric and machine learning models might be explained by the way they capture prediction patterns from the data. The machine learning models prioritise capturing nonlinear prediction patterns (Ryo and Rillig 2017). The lower prediction performance of machine learning models for the DTA trait may result from their inability to construct simpler models focusing on linear effects, leading to overfitting (Hawkins 2004). Hence, the rrBLUP and BayesB, focused on capturing linear patterns, outperformed the machine learning models.

In contrast, the TILN trait, with a higher potential to exhibit nonlinear patterns, the performance may have been a more suitable prediction problem for the machine learning models. However, the machine learning models may not have been able to capture all interactions in the complex network using a small amount of training data, resulting in similar prediction performances to the parametric genomic prediction models. The different prediction performance of SVR across the two traits could be an example of this. Using the kernel of the radial basis function, SVR mainly targeted complex nonlinear prediction patterns, which may have generated models of too much complexity to accurately predict the DTA trait. For the TILN trait, this kernel could have enabled SVR to capture some of the complex interactions. These observations indicate that complexity of gene and trait networks underlying target traits in crop breeding programs can be a crucial factor affecting the performance of genomic prediction models.

### No consistent winner amongst individual genomic prediction models

The lack of a consistent winner among the individual genomic prediction models (Figure 3 and S1) poses the relevance of the No Free Lunch Theorem (Wolpert and Macready 1997) for genomic prediction problems. For the DTA trait, while rrBLUP and BayesB slightly outperformed the other models by rrBLUP and BayesB was observed, all the individual genomic predictions except SVR, showed similar prediction accuracy and error. This absence of a best individual genomic prediction model was also evident in the TILN trait, with no clear differences in prediction accuracy and error among the models. The No Free Lunch Theorem is further supported by the positive associations between predicted phenotypes from different genomic prediction models (Figure 4). Each individual genomic prediction model returned similar predicted phenotypes for the same RILs, resulting in similar prediction performance. Therefore, the distinctive algorithms of the individual genomic prediction models did not result in significantly diverse prediction performances.

The lack of a consistent winner among individual models has been observed for prediction problems in other fields. For instance, Fernández-Delgado *et al*. (2014) tested 179 prediction models over 121 datasets and concluded that RF achieved the overall highest prediction performance. However, no statistical performance superiority of RF to the second best (support vector machine leveraging Gaussian kernel) was detected in their experiment, indicating a subtle performance difference. The results from Gómez and Rojas (2016) also showed that no individual model clearly outperformed the others. The absence of a prediction model that was immune to all the negative factors (noise from datasets, data imbalance and dissatisfaction with model assumptions) was considered to be the cause of finding no single “best” individual model. Similarly, other research (Merrick and Carter 2021; Plavšin *et al*. 2022) suggested the difficulty in finding a single “best” individual model. These results indicate that some prediction individual models can outperform others in specific scenarios but are not universally superior.

Therefore, we argue that focusing on developing an individual genomic prediction model for diverse tasks is not strategic. Instead, leveraging the expectations from the Diversity Prediction and No Free Lunch Theorems, ensemble approaches can be one solution to overcome the limitations of optimising prediction-based crop breeding around individual genomic prediction algorithms.

### Ensemble improved prediction performance

One clear consensus, from the results of this study, is the improved predictive ability of the naïve ensemble-average model compared to the individual genomic prediction models. For the DTA and TILN traits, the median prediction accuracy of the naïve ensemble-average model was the highest across the RIL populations. Although the naïve ensemble-average model did not achieve the lowest MSE for the DTA trait, it was almost equivalent to rrBLUP and BayesB that achieved the lowest MSE (Figure 3 and Table 2). When compared against the average of the individual genomic prediction models, the naïve ensemble-average model consistently outperformed the individual genomic prediction models on the basis of prediction accuracy and error (Figure 2). These results suggest an opportunity to improve prediction performance with the naïve ensemble-average model.

The success of the naïve ensemble-average model is derived from the diversity of information contributed by multiple, individual genomic prediction models. The range of associations among the estimated genomic marker effects of the individual genomic prediction models (Figure 4) illustrates this diversity These ranges of associations are a result of algorithmic differences that differentially weight predictive features from the same input data. This phenomenon, called the Rashomon effect (Breiman 2001b), states that the sets of models (Rashomon sets) capture different effects of features from the same datasets due to distinct properties of the prediction algorithms. Ensemble models can use this prediction diversity to generate a more comprehensive representation of the prediction problem. In the case of this study, providing a more comprehensive view of trait genetic architecture. Hence, prediction diversity from multiple, individual genomic prediction models is a core factor behind the improved prediction performance of the naïve ensemble-average model.

Although the set of the six individual genomic prediction models chosen is one of many possibilities, their contrasting algorithms for estimating genetic effects contributed to the diversity in effect estimates. For example, the parametric models (rrBLUP and BayesB) primarily target capturing SNP main effects, while the semi-parametric model RKHS consider SNP interaction terms in addition to SNP main effects. Nonparametric machine learning models prioritise SNP interactions by considering nonlinear relationships between SNPs. Each distinctive prediction algorithm develops a unique hypothetical space, and the true value can be outside the space of a particular individual genomic prediction model. Creating a new hypothetical space through ensembles of individual models can provide predictions outside the hypothetical spaces of any one, individual model. (Johnson and Giraud-Carrier 2019; Dietterich 2000). Complementing the respective errors of each genomic prediction model in the ensemble enables the construction of solutions for complex tasks with higher performance (Dong *et al*.2020; Kick and Washburn 2023).

The impact of information diversity from individual genomic prediction models was also elucidated in terms of the Diversity Prediction Theorem (Page 2018). As shown in Table 2, prediction diversity was observed among the individual genomic prediction models, contributed to the reduction of the ensemble error. Consequently, the prediction accuracy and error of the naïve ensemble-average model was higher for both traits. Thus, the level of information diversity among the individual prediction models critically influenced the performance of the naïve ensemble-average model from a theoretical view as well.

More broadly, the higher predictive ability of the naïve ensemble-average model could also be viewed as a result of avoiding stagnation in local optima. Each of the six genomic prediction modelling algorithms may achieve different local optima within their possible prediction space, the size of which depends on the given data and algorithms. Algorithms trapped in their local optima can inhibit opportunities to explore the broader problem state space closer to the global optima, where more precise predictions may be achieved. Applying another algorithm on top of a set of individual genomic prediction models can increase the likelihood of shifting from the local to the global optima. For example, Wu *et al*. (2019) discussed the benefits of using an ensemble concept in population-based optimization approaches (a method containing a number of adaptive prediction models to find global optima), suggesting that the global optima can be efficiently discovered by cooperatively sharing information from each prediction model rather than using them independently. The advantage of discovering global optima by an ensemble has also been mentioned in evolutionary algorithms (iterative model improvement adaptively done with observed prediction results) (Yu and Suganthan 2010) and metamodeling (representation of a model to a simpler mechanism compared to the original one) (Ferreira and Serpa 2018) as well. In short, the ensemble can help reach global optima using information derived from different local optima.

### Future Opportunities

Several components can be considered to improve the prediction performance of the genomic prediction models considered in our investigation; hyperparameter tuning and weight optimization. Below, we briefly discuss each of these components.

In this experiment, hyperparameters of the genomic prediction models have been tuned heuristically rather than systematically. Systematic hyperparameter tuning can be conducted by approaches such as cross-validation and Bayesian methods. Prior studies (Sandhu *et al*. 2021b; Kick *et al*. 2023) have demonstrated that hyperparameter tuning can increase prediction performance. Due to the computational limitations and the data size (the total number of RILs), the hyperparameter tuning was not implemented and default hyperparameter values were primarily used. Hence, optimising hyperparameters may improve prediction performance for both individual and ensemble genomic prediction models by overcoming these computational and data limitations.

Lastly, performance improvements can be expected by optimizing the weights applied to the predicted phenotypes in ensembles of genomic prediction models. In this experiment, the predicted phenotypes were “naïvely” averaged from the six individual genomic prediction models by assigning equal weight to each. In other words, the six individual genomic prediction models contributed equally to the ensemble averaging step. However, this is one of many possible weighting approaches. In some applications, higher prediction performance was achieved by tuning weights based on the prediction performance of each individual model (Liang *et al*. 2021; McCormick *et al*. 2021; Yu *et al*. 2021; Wang *et al*. 2023). Model selection is an extreme scenario where the weight of some individual models is set to zero based on their contribution level, improving prediction performance of ensembles in in several cases (Zhou *et al*. 2002; Li *et al*. 2004; Huang and Wei 2022). Therefore, weight optimization could provide further improvements tin the prediction performance of ensembles of many models.

## Conclusion

We investigated the prediction performance of the naïve ensemble-average model compared to six individual genomic prediction models for the DTA and TILN traits in the TeoNAM dataset. Our results showed higher prediction accuracies and lower prediction errors with the ensemble model compared to individual genomic prediction models. Therefore, ensemble approaches could be a promising tool for genomic prediction. The increased predictive ability of the ensemble model is derived from the diversity of prediction outcomes among the individual genomic prediction models, as explained by the Diversity Prediction Theorem. Further research is needed to investigate the effectiveness of ensemble approaches on other datasets. Our results suggest that the ensemble can improve selection accuracy and reduce prediction errors, demonstrating the potential to accelerate genetic gain in breeding programs.

## Data and code availability

All the datasets leveraged in this study were collected by Chen *et al*. (2019) and full RILs and phenotype data are publicly available at panzea/genotypes/GBS/TeosinteNAM and Supplemental_Material_for_Chen_et_al_2019/9250682 respectively. The code generated for this experiment is shared at https://github.com/ShunichiroT/ensemble. The data and code used in this study were also uploaded at https://zenodo.org/records/14776591.

## Acknowledgments

We thank the National Computing Infrastructure (NCI) and Research Computing Centre (RCC) at the University of Queensland for supporting our experiment through High Performance Computing (HPC) machines.

