## Supplementary Material for "Improved Genomic Prediction Performance with Ensembles of Diverse Models"

**Supplementary Material 1: Other Supporting Results**

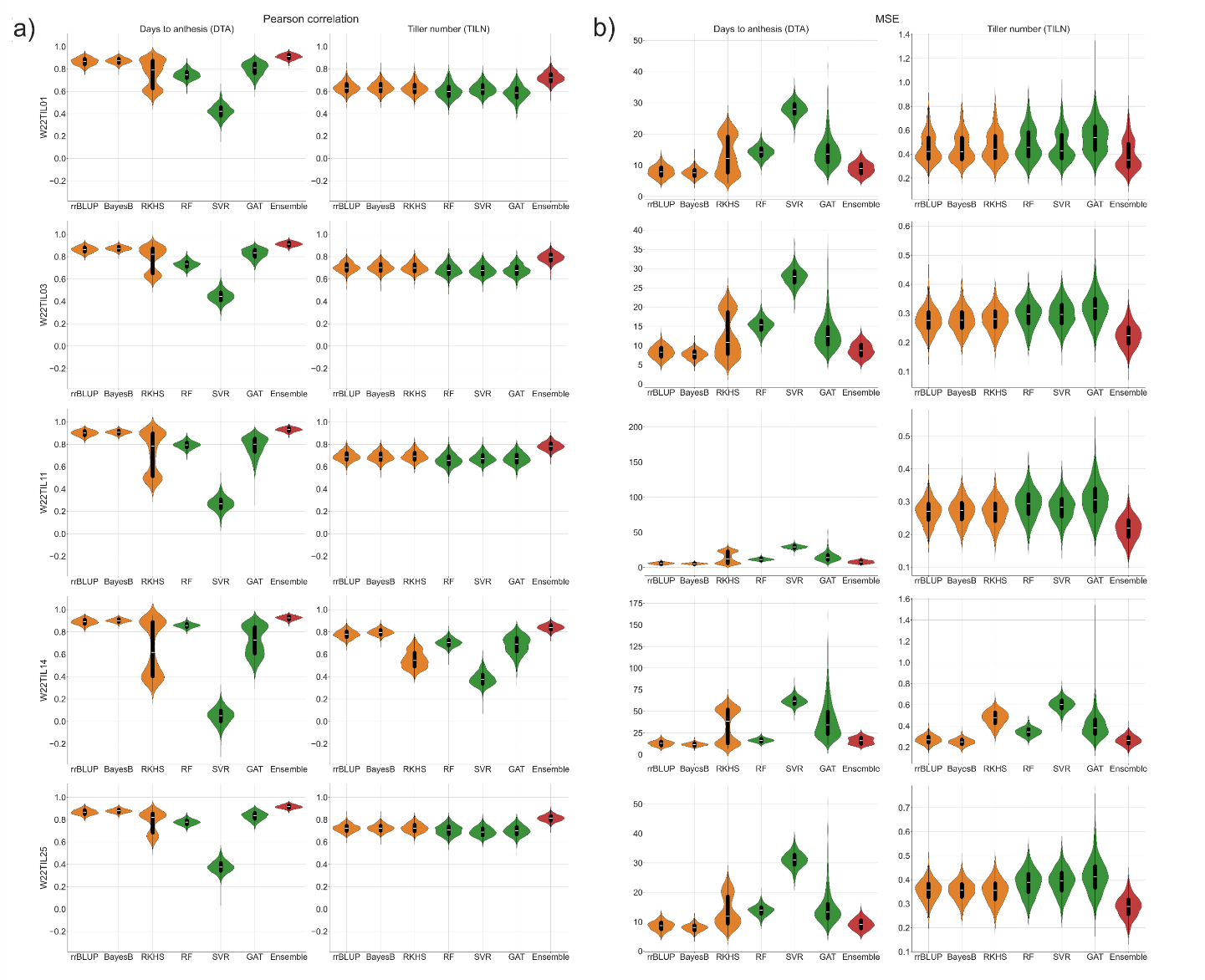

**Figure S1** A comparison of genomic prediction performance of the naïve ensemble-average model (Ensemble) versus each of the individual genomic prediction models at the per-population level in violin plots. The width of the violins represents the distribution of metric values for predictions from all combinations of the five RIL populations, three training-test ratios and 500 random samples. The performance of genomic prediction models was measured with a) the Pearson correlation and b) mean squared error (MSE). The orange represents the performance of classical models (rrBLUP, BayesB and RKHS) while the green represents machine learning models (RF, SVR and GAT). The red is the performance of the ensemble (naïve ensemble-average). Box plots within the violin plots represent the median metric value (white line) and the interquartile range (black box) with whiskers extending 1.5 past the interquartile range.

**Table S1 Pairwise correlation of individual genomic prediction models at the level of predicted phenotypes (the top right of the diagonal) and genomic marker effects (the bottom left of the diagonal) in the DTA and TILN traits, corresponding to the scatter plot matrices in Figure 4.**

|  | DTA | | | | | | TILN | | | | | |
| --- | --- | --- | --- | --- | --- | --- | --- | --- | --- | --- | --- | --- |
|  | rrBLUP | BayesB | RKHS | RF | SVR | GAT | rrBLUP | BayesB | RKHS | RF | SVR | GAT |
| rrBLUP | 1.000 | 0.976 | 0.804 | 0.746 | 0.362 | 0.673 | 1.000 | 0.991 | 0.944 | 0.749 | 0.718 | 0.714 |
| BayesB | 0.742 | 1.000 | 0.759 | 0.770 | 0.355 | 0.676 | 0.855 | 1.000 | 0.924 | 0.752 | 0.710 | 0.711 |
| RKHS | 0.022 | 0.022 | 1.000 | 0.584 | 0.475 | 0.602 | 0.081 | 0.083 | 1.000 | 0.743 | 0.748 | 0.706 |
| RF | 0.351 | 0.622 | 0.019 | 1.000 | 0.311 | 0.635 | 0.383 | 0.536 | 0.035 | 1.000 | 0.869 | 0.854 |
| SVR | 0.084 | 0.197 | -0.021 | 0.274 | 1.000 | 0.478 | 0.480 | 0.514 | 0.005 | 0.445 | 1.000 | 0.868 |
| GAT | 0.207 | 0.277 | 0.017 | 0.280 | 0.237 | 1.000 | 0.320 | 0.351 | 0.035 | 0.278 | 0.383 | 1.000 |

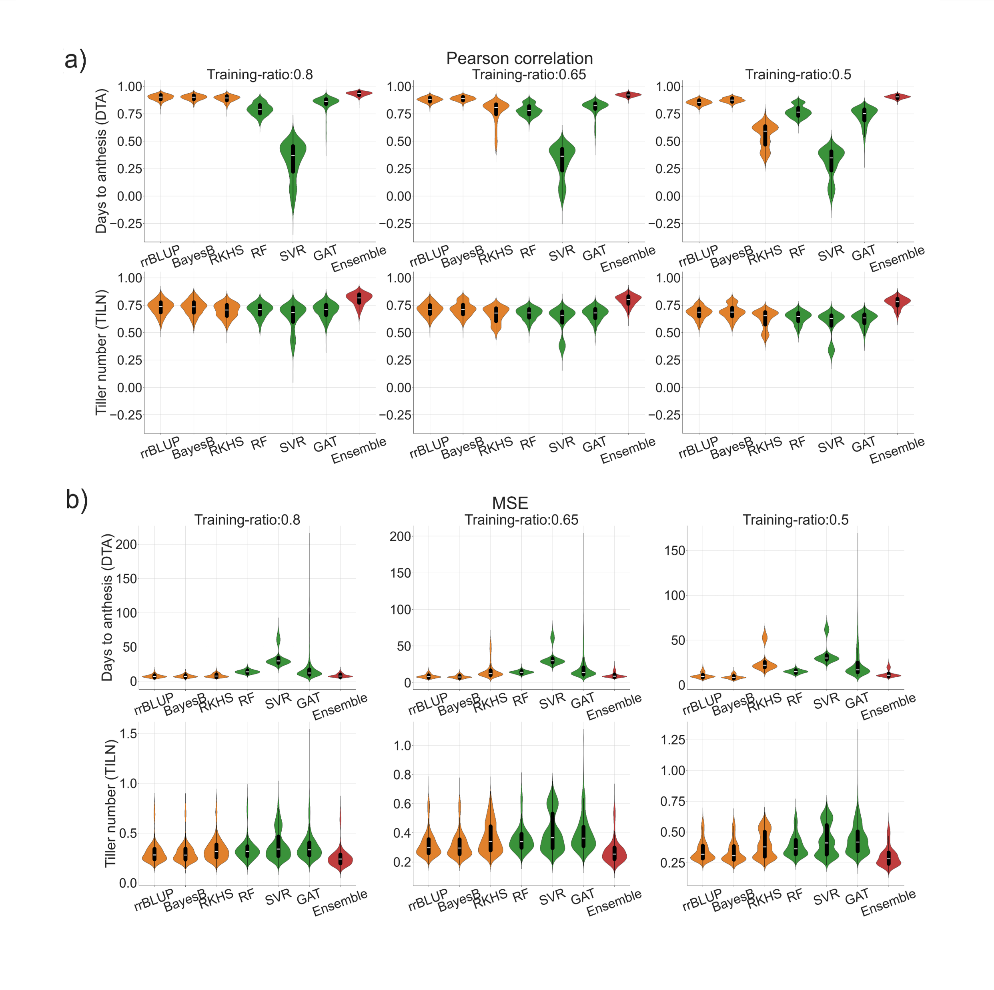

**Figure S2** A comparison of genomic prediction performance of the naïve ensemble-average model (Ensemble) versus each of the individual genomic prediction models per training-test ratio across RIL populations in violin plots. The width of the violins represents the distribution of metric values for predictions from all combinations of the five RIL populations, three training-test ratios and 500 random samples. The performance of genomic prediction models was measured with a) the Pearson correlation and b) mean squared error (MSE). The orange represents the performance of classical models (rrBLUP, BayesB and RKHS) while the green represents machine learning models (RF, SVR and GAT). The red is the performance of the ensemble (naïve ensemble-average). Box plots within the violin plots represent the median metric value (white line) and the interquartile range (black box) with whiskers extending 1.5 times the interquartile range.

**Supplementary Material 2: Investigating an Alternative Imputation Method**

All analyses reported in this section were conducted on the TeoNAM data set following the imputation of missing markers using flanking markers. When suitable flanking markers were available and both flanking markers possessed the same parental allele phase, markers with missing calls were imputed with the parental allele phase of their flanking markers. If flanking markers on both sides contained different parental allele phases, the parental allele phase of the closest flanking marker was leveraged to impute the missing marker calls. When SNPs contained more than 10% of missing marker calls, the imputation approach was not feasible, and these SNPs were removed from the data set. Similarly, if the allele calls of the entire set of markers in a chromosome were missing, the corresponding RILs were removed from the data set.

When the target trait phenotype for an RIL was missing the RIL without the target trait phenotypes was removed from the data set.

Applying the equation for the Diversity Prediction Theorem the value for each term in the equation was calculated per prediction scenario (Experimental Flow) to evaluate the ensemble error with the consideration of the diversity of prediction models. After the calculation, the value for each term was averaged at per trait level.

**Results**

### **High diversity in the predicted phenotypes of models promoted the reduction of ensemble error**

The ensemble error with higher predicted phenotype diversity from the individual genomic prediction models improved the ensemble error in the Diversity Prediction Theorem (Table 2, S2). The diversity of prediction model prediction for the most frequent allele imputation (7.09 for DTA and 0.09 for TILN) was higher than the alternative (flanking marker) imputation (6.16 for DTA and 0.04 for TILN) for both DTA and TILN traits. An increase in the diversity of model predictions leads to the reduction of the ensemble error. Hence, the estimated ensemble error for the most frequent allele imputation approach (10.17 for DTA and 0.28 for TILN) was lower than for the alternative flanking marker imputation approach (10.50 for DTA and 0.32 for TILN). The higher prediction performance for the former was observed in higher prediction accuracy and lower prediction error of the naïve ensemble-average model compared to the latter imputation approach (Figure 2, 3, S3, S4). The diversity in individual genomic prediction models was also observed by showing fewer positive associations at both predicted phenotypes and genomic marker effects for the most frequent allele imputation approach (Figure 4) compared to the alternative flanking marker imputation approach (Figure S5). The comparison of the prediction result from the two imputation approaches emphasizes that higher information diversity from the individual genomic prediction models led to prediction performance improvement by the naïve ensemble-average model for both imputation methods.

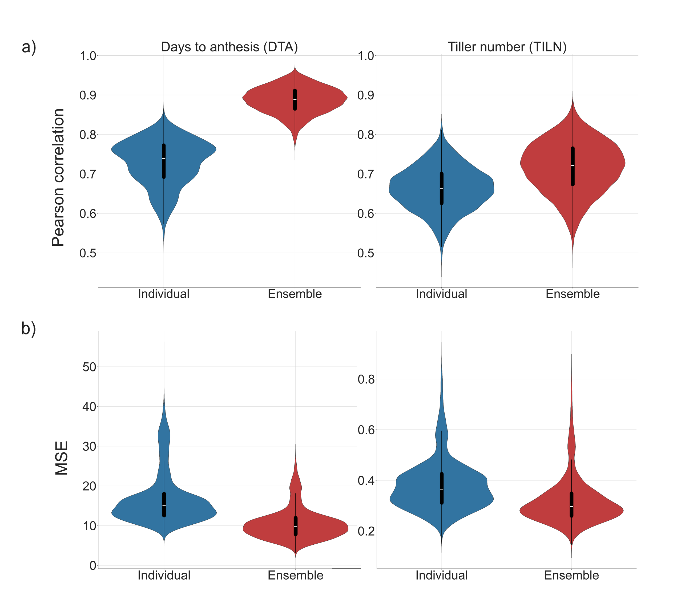

**Figure S3** Violin plots comparing genomic prediction performance of the average of individual genomic prediction models (Individual) in the blue versus the naïve ensemble-average model (Ensemble) in the red for the alternative (flanking marker) imputation approach. The performance of genomic prediction models was measured with a) the Pearson correlation and b) mean squared error (MSE). The width of the violins represents the distribution of metric values for predictions from all combinations of the five RIL populations, three training-test ratios and 500 random samples. Box plots within the violin plots represent the median metric value (white line) and the interquartile range (black box) with whiskers extending 1.5 times the interquartile range.

**Table S2 Estimates of the average value for each term of the Diversity Prediction Theorem (Equation (1)) across 7,500 scenarios in the alternative (flanking marker) imputation approach for the traits days to anthesis (DTA) and tiller number per plant (TILN)**

|  | ensemble error  (first term) | | average error  (second term) | | prediction diversity  (third term) |
| --- | --- | --- | --- | --- | --- |
| DTA | 10.50±1.43 | 16.66±2.09 | | 6.16±0.95 | |
| TILN | 0.32±0.05 | 0.36±0.06 | | 0.04±0.00 | |

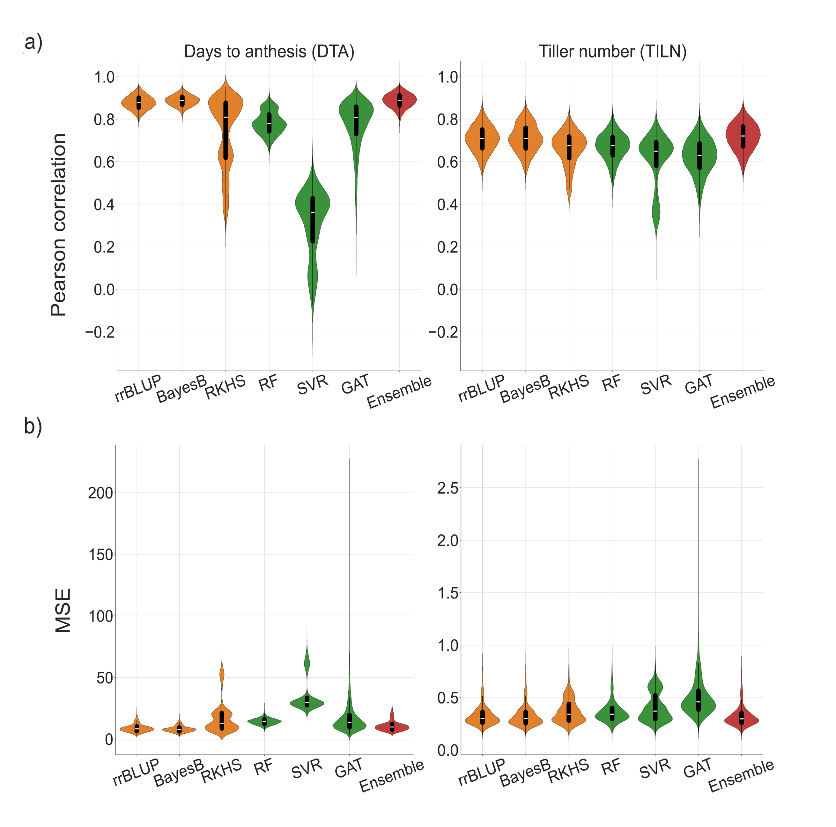

**Figure S4** A comparison of genomic prediction performance of the ensemble (naïve ensemble-average) model versus each of the individual genomic prediction models in violin plots for the alternative (flanking marker) imputation approach. The width of the violins indicates the distribution of the metric values for predictions from all combinations of the five RIL populations, three training-test ratios and 500 random samples. The performance of genomic prediction models was measured with a) the Pearson correlation and b) mean squared error (MSE). The orange represents the performance of classical models (rrBLUP, BayesB and RKHS) while the green represents machine learning models (RF, SVR and GAT). The red is the performance of the ensemble. Box plots within the violin plots represent the median metric value (white line) and the interquartile range (black box) with whiskers extending 1.5 times the interquartile range.

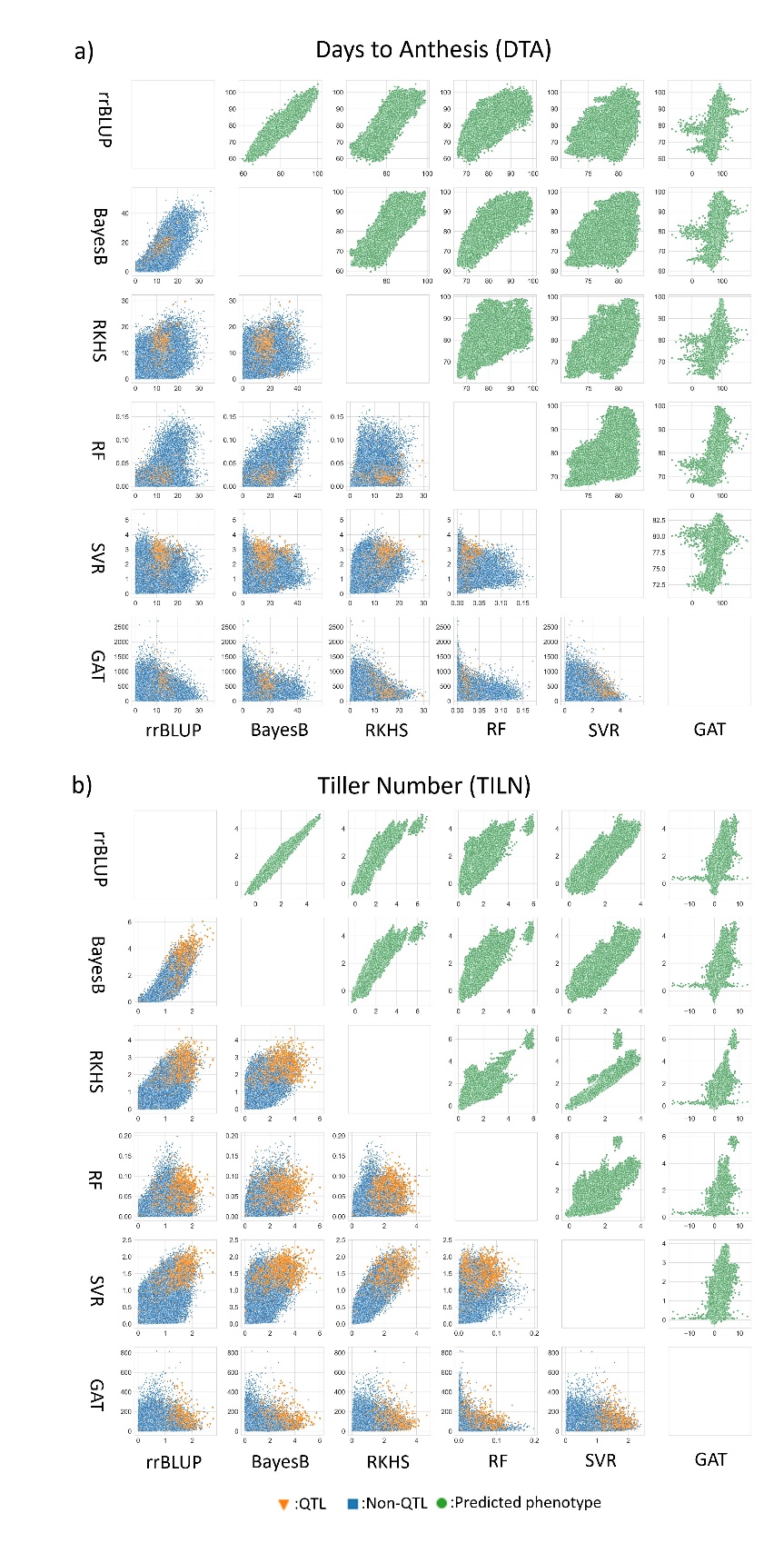

**Figure S5** Pairwise comparison of individual genomic prediction models at predicted phenotypes (top right triangle) and genomic marker effects (the bottom left triangle) levels for both traits; a) the days to anthesis (DTA) and b) the tiller number per plant (TILN) for the alternative (flanking marker) imputation approach. The green circle dots represent a pair of predicted phenotypes for RILs included in the test set in each sample scenario. The blue square and orange triangle dots indicate a pair of estimated genomic marker effects in each sample scenario classified as non-QTL and QTL by Chen *et al.* (2019) respectively. A genetic marker was classified as a QTL if it was the closest to a QTL position within the support interval of 2 logarithm of the odds (LOD) calculated by Chen *et al.* (2019). Each point represents predicted phenotypes or genomic marker effect of each SNP in predictions from all combinations of the five RIL populations, three training-test ratios and 500 random samples.
